# Unraveling candidate genomic regions responsible for delayed post-harvest deterioration in Cassava (*Manihot esculenta* Crantz)

**DOI:** 10.64898/2026.06.11.731395

**Authors:** Diana C Solarte Certuche, Gabriel M de Freitas, Jean-Luc Jannink, Cinara Fernanda Garcia Morales, Tamires Sousa Cerqueira, Bruna Santos de Santana, Eder Jorge de Oliveira, Antônio Augusto F Garcia

**Affiliations:** São Paulo University, Genetics Department, Piracicaba, São Paulo, Brazil; USDA-ARS, Robert W. Holley Center, Ithaca, NY, United States of America; Embrapa Mandioca e Fruticulture, EMBRAPA, Genetic Resources and Cultivar Development Center, Cruz das almas, Bahia, Brazil; Federal Unviersity of Recôncavo da Bahia, CCAAB, Cruz das Almas, Bahia, Brazil

**Keywords:** Cassava, GWAS, Post-harvest physiological deterioration

## Abstract

Post-harvest physiological deterioration (PPD) represents a significant challenge of cassava production and commercialization. This multifaceted biological process involves a series of mechanisms, including enzymatic stress responses, alterations in gene expression, protein synthesis, accumulation of secondary metabolites, and ultimately, programmed cell death. These changes render the storage roots unpalatable and unmarketable. Therefore, unraveling the genetic architecture of PPD and exploring the interactions of associated genes during its early and late stages is essential for the crop production. We used modern genetic resources to unravel the genetic basis of PPD, based on a genome-wide association study (GWAS), utilizing a combination of different models, including BLINK (Bayesian-information and Linkage-disequilibrium Iteratively Nested Keyway), SUPER (Settlement of MLM Under Progressively Exclusive Relationship), and MLMM (Multi-locus mixed models). The phenotyping dataset spanned five years and included evaluations from 42 different trials, evaluating the Embrapa (Brazilian Agricultural Research Corporation) germplasm along with a population derivative from a genomic selection cycle. We utilized a genotype dataset comprising 26,000 high-quality SNPs (single nucleotide polymorphisms). Our findings indicated four significant genetic variants located on chromosomes 2, 5, and 13, which together explain 35.83 % of the phenotypic variation. These variants are associated with genes that are closely linked to the pathways activated during the early and late symptoms of PPD. The identification of these three key genes provides valuable insights into the genetic architecture of PPD and lays a strong foundation for molecular breeding, supporting the efforts to identify cassava genotypes with enhanced PPD tolerance, the identified genomic regions may be incorporated into genomic selection models, thereby enhancing marker-assisted selection (MAS) and improving breeding strategies for long shelf life and high-quality agronomic cassava cultivars for the cassava community.

## Introduction

Cassava (*Manihot esculenta* Crantz) has become a globally significant crop, with global production increasing by approximately 16–17% over the past decade. By 2024, global production exceeded 315 million tonnes, with Nigeria remaining the largest producer (∼62.69 million tonnes) and Brazil ranking fifth (∼18.51 million tons) (FAO, 2024). Cassava’s agronomic resilience, including broad edaphic adaptability and notable drought tolerance, makes it a critical crop for both smallholder and commercial farming systems (Borku et al., 2025). Historically, cassava has underpinned food security in many developing countries, particularly in Africa, where it is the third most important staple food. At the same time, the demand for cassava-derived products has expanded across an increasingly diverse set of industries (food, beverages, confectionery, textiles, paper-making, pharmaceuticals, cosmetics, chemicals, and bioethanol) (Mohidin et al., 2023). This increase in demand has sparked significant scientific interest, as evidenced by the 21,136 related publications focused on cassava in the Scopus database (Otekunrin, 2024).

Despite its agronomic advantages, cassava breeding and commercialization face several persistent constraints embedded in the base germplasm: low yield potential in some clones, susceptibility to major pests and diseases, low micronutrient content in many cultivars, and elevated cyanogenic potential in others. Among these limitations, post-harvest physiological deterioration (PPD) of storage roots poses a particularly acute challenge for transport, storage and marketability (Devi et al., 2022). PPD is a complex, wound-triggered physiological syndrome that initiates in root parenchyma tissue after harvest. Mechanical damage during harvest activates a rapid enzymatic stress response that propagates signaling from wounded zones to distal tissues and triggers defense-related gene expression (Saravanan et al., 2016). The organoleptic and nutritional consequences of PPD, principally vascular parenchyma discoloration and loss of starch and other quality attributes, significantly reduce commercial value and consumer acceptance (Devi et al., 2022). Global production losses due to PPD have been estimated at ∼19 %, with losses in regions having poor limited transport and storage infrastructure (e.g., many areas in Africa) reaching ∼29 % (Saravanan et al., 2016). Extending the shelf life of cassava roots during storage is a common goal. This not only helps address food security challenges but also enhances marketing opportunities and facilitates the large-scale conversion of cassava into a commercial crop.

At the cellular and biochemical levels, PPD progresses through well-documented stages. Early responses include extensive reprogramming of gene expression, altered protein synthesis and accumulation of secondary metabolites (Iyer et al., 2010). An oxidative burst, marked by production of superoxide radicals and other reactive oxygen species (ROS), is a hallmark early event; ROS together with hydrogen peroxide modulate transcriptional networks and secondary metabolism (Lebot et al., 2023; Reilly et al., 2007). These redox dynamics are accompanied by increased respiration and conversion of starch to soluble sugars, producing a measurable decline in starch content during storage (Zainuddin et al., 2018). Concomitantly, volatile metabolites (cyanohydrins, aldehydes, ketones, alcohols) and ethylene are produced (Shi Liu et al., 2017), and Ca2+ signaling appears to contribute to the induction of oxidative bursts (Salcedo, 2011). Oxidation of hydroxycoumarins such as scopoletin by ROS and peroxidases yields colored complexes that explain the characteristic blue–dark discoloration observed in affected roots (Shi Liu et al., 2017). Recent evidence further indicates that phenolics and carotenoids can modulate damage to metabolic pathways and thereby mitigate symptom severity in early PPD stages (Che et al., 2024). Later stages progress to programmed cell death, secondary microbial colonization, and tissue softening, culminating in severe quality losses (Zainuddin et al., 2018).

Overall, PPD is a complex phenomenon that results from interactions between different pathways, influencing whether a genotype shows tolerance or susceptibility to PPD. Given this substantial economic and food-security impact, extensive biochemical, molecular and genetic studies have sought to elucidate PPD mechanisms and identify tolerant germplasm (Buschmann et al., 2000; Iyer et al., 2010; Djabou et al., 2017; Lebot et al., 2023; Che et al., 2024; Cortes et al., 2002; Reilly et al., 2007; Ma et al., 2023; Ayalde et al., 2024). Breeding work has also explored phenotypic sources of tolerance and estimated genetic parameters for PPD (Kawano & Rojanaridpiched, 1983; Aina et al., 2007; Akinwale et al., 2010; Tumuhimbise et al., 2015; Venturini et al., 2016). Although significant knowledge has been gained regarding PPD in cassava, achieving long shelf-life genotypes remains a major challenge for breeding programs and stakeholders across the cassava value chain. PPD is notably difficult to breed for due to the strong influence of environmental conditions on trait expression. Developing longer shelf-life genotypes involves navigating a complex genetic structure, where numerous small genetic effects contribute to the polygenic nature of PPD symptom expression. Additionally, there is a negative correlation between PPD and key traits for cassava, such as dry matter content and carotenoids. Over the past few decades, considerable efforts have been made to clarify this intricate genetic structure, including the identification of several quantitative trait loci (QTL) in mapping populations in African germplasm. However, these QTL analyses have not accounted for a significant portion of the phenotypic variation observed. Despite extensive research, to our knowledge there have been no successful applications of marker-assisted breeding strategies to improve postharvest traits related to PPD reported to date.

Markers identified in association with PPD could be utilized as covariates in genomic selection (GS) and marker-assisted selection (MAS) as routine practices in the breeding pipeline. This would provide a viable solution to obtain long shelf life post-harvest accessions while also addressing other important agronomic traits in different cassava varieties, such as dry matter content, cyanogenic compounds, carotenoids, and resistance to major diseases (Mbinda & Mukami, 2022).

In this study, we build upon previous efforts to understand the genetic architecture of PPD by integrating multi-environment and multi-year phenotyping with an appropriate association mapping strategy. This approach enables us to better control for environmental interactions affecting PPD expression. As a result, we provide a comprehensive catalog of trait-locus combinations and candidate genes that can explain the observed variation. Building on prior statistical analyses of Brazilian germplasm that revealed substantial variability and environmental effects on PPD expression (Venturini et al., 2016), the present study investigates the genetic architecture underlying PPD in a broad Brazilian germplasm collection. We extend Venturini et al.’s work by including a new population derived from recent genomic selection cycles in the Embrapa cassava breeding program. Using genome-wide association studies (GWAS) implemented with multiple statistical models to account for population structure and kinship, we analyzed 812 cassava clones genotyped by Genotyping-by-Sequencing (GBS) and scored with 27,047 SNPs. These markers obtained could be utilized as covariates in (GS) and (MAS) as steps routine practices in the breeding pipeline. This would provide a viable solution to obtain long shelf-life post-harvest accessions.

## Material and Methods

### Plant material and field trials

The study evaluated a panel of 812 cassava clones derived from the Embrapa cassava breeding program (Supplementary Table 1). Phenotypic data were collected across 42 field trials conducted in multiple years and locations throughout the state of Bahia, Brazil (Supplementary Table S2). The trials represent historical and ongoing breeding evaluations that include germplasm collections and populations derived from four genomic selection cycles (C0, C1, C2, and C3). Experimental designs varied by year and location as follows: in 2018 germplasm and the C0 population were evaluated at a single location using an augmented block design; in 2020 evaluations were expanded to 10 locations using a combination of augmented block and randomized complete block designs (RCBD); in 2021 the panel was assessed across 13 locations using an augmented block design, RCBD and a completely randomized design (CRD); 2022 trials included two designs across 10 locations; and 2023 trials covered eight locations. Plot-level management, planting density and standard crop husbandry followed local Embrapa protocols for cassava (Filho, 2013).

### Post-harvest physiological deterioration (PPD) phenotyping

PPD was assessed using two complementary protocols to quantify root deterioration: (i) a conventional visual scoring of the proportion of damaged root tissue on a 0–100 scale (Venturini et al., 2016), and (ii) an image-based measure obtained from digital photographs processed in *ImageJ software v.2.14.0* (Schneider et al., 2012), employing automated area-detection routines and machine-learning plugins to estimate the percentage of affected tissue (scaled 0–100). For both methods, freshly harvested roots were cleaned and sanitized, then each root was cut into six longitudinal slices. Slices were stored for 10 days on open shelves beneath a metal-roofed shelter that allowed free air circulation on all sides. After the storage period, slices were photographed following a standardized imaging protocol (fixed camera-to-sample distance, consistent lighting, inclusion of a scale bar and color reference), and the resulting images were evaluated using both the visual scoring and ImageJ pipelines.

### DNA extraction and SNP genotyping

DNA extraction and SNP genotyping were conducted by collaborators from Embrapa as part of a collaborative effort, following the protocol described below. Genomic DNA was isolated from young, freeze-dried leaf tissue using the cetyltrimethylammonium bromide (CTAB) extraction method by Doyle & Doyle, 1987. Genome-wide genotyping was performed using GBS following standard protocols; raw sequence reads were aligned to the cassava reference genome (v6) and variant calling was performed with the *Genome Analysis Toolkit (GATK v. 4.4)* pipeline (McKenna et al., 2010). Initial quality control removed markers with minor allele frequency (MAF) < 0.01 and sites with >60 % missing genotype calls, producing an initial high-quality set of 27,047 biallelic SNPs for downstream filtering and analyses. All genotype matrices were quality-checked and formatted for population-structure, kinship and association analyses.

### Phenotypic data analysis

Phenotypic data used in this study were provided by collaborators from Embrapa and were collected according to the Embrapa’s protocol. Phenotypic traits were analyzed using a linear mixed model using the package *ASReml-R v.4.1* (Butler et al., 2023) implementing a two-stage mixed-model framework adapted for multi-environment trial data (Endelman, 2023). In the first stage, environment-specific linear mixed models were fitted to plot-level PPD observations to adjust for design effects and to obtain genotype adjusted means Best Linear Unbiased Estimator (BLUEs) within each environment. The general form of the per-environment statistical model was:

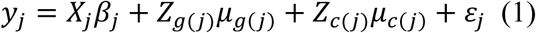

where, ***y*_j_** represents the data vector of PPD evaluation plot in the location-year combination that represents the environment. ***X***_***j***_ is an incidence matrix relating observations in enviroment j, ***β***_***j***_ is a vector that represents the fixed effects specific for environment j^th^, ***Z***_***g***(***j***)_ is an incidence matrix for clones, ***μ***_***g***(***j***)_ is a vector that contains the random effects of the clone in environment j^th^, ***Z***_***c***(***j***)_ incidence matrix for additional random design- related effects in environment j^th^, such as row and columns, and blocks, ***μ***_***c***(***j***)_ vector of random effects between environments in the combination location-year, and, ***ε***_***j***_ vector of plot error associated with the environment j^th^. Assumed that ***ε***_***j***_ ∼ 𝓝(**0**, ***g***_***j***_), ***μ***_***g***(***j***)_ ∼ 𝓝(**0**, ***g***_***j***_), ***μ***_***c***(***j***)_ ∼ 𝓝(**0**, ***c***_***j***_).

In the second stage, environment-level genotype estimates were combined to obtain single genotype genetic values. De-regressed BLUPs (Best Linear Unbiased Prediction) (or weighted BLUPs) were computed to account for heterogeneous error variances among environments. The second stage working model can be expressed as:

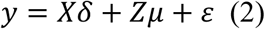

where ***y*** is the vector of de-regressed genotype values, ***X*** is the design matrix relating genotype level observations to fixed effects across environments, ***δ*** contains fixed effects (environmental, markers and inbreeding), ***Z*** is the incidence matrix relating observations to genotypes, ***μ*** represents the vector of random genetic effects structured by a marker-based relationship matrix, and ***ε*** is the vector of residuals.

The resulting BLUPs values obtained in this analysis were used to the GWAS analysis model in the next steps.

Broad sense heritability for the trait was estimated on an entry mean basis using the equation

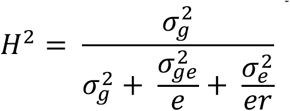

Where 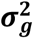 is the genotypic variance, 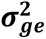 is the genotype by environment interactions variance, 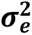 error variance, and respectively r and e denote the number of replications and the number of environments included in the analysis, respectively.

### Population structure, kinship and linkage disequilibrium

Population structure was evaluated using multiple complementary approaches. Bayesian clustering was performed with *STRUCTURE v2.3.4* (Pritchard et al., 2000) with multiple independent runs across K=1 to K=6; runs included appropriate burn-in and Markov Chain Monte Carlo (MCMC) sampling and were replicated to ensure convergence. The most likely number of clusters (*Δ*K) was determined with *STRUCTURE HARVESTER v.0.6.9.94* (Earl & vonHoldt, 2012) using the method of Evanno et al. (2005). Principal component analysis (PCA) was performed in *TASSEL software v. 5.0* (Bradbury et al., 2007). Pairwise linkage disequilibrium (LD) between markers was computed using *PLINK v. 2.0* (Purcell et al., 2007) and LD decay and local r² patterns were evaluated to define candidate regions, considering SNP pairs with r² ≥ 0.8 as being in strong LD. Kinship among individuals was estimated using a marker-based genomic relationship matrix was calculated following the method of Theo H. E. VanRaden (2008), based on centered genotype data and allele frequencies, where higher values indicate greater genetic similarity. Visualization of structure and clustering results was generated with construct (Bradburd et al., 2016) and standard R plotting tools.

### Genome-wide association study (GWAS)

The association analyses used BLUPs derived from 812 clones evaluated across multiple location–year combinations as the response variable. Genotype data represented in 27,044 high-quality, biallelic SNPs. GWAS was performed in *GAPIT v3* (Wang & Zhang, 2021) using several complementary multi-locus and mixed-model methods to improve power and reduce false positives: *i*) SUPER (Settlement of MLM Under Progressively Exclusive Relationship) (Wang et al., 2014), *ii*) BLINK (Bayesian-information and Linkage-disequilibrium Iteratively Nested Keyway), which employs linkage disequilibrium to streamline marker selection (Huang et al., 2019), and *iii*) MLMM (Multi-locus Mixed Model), which adjusts for cofactors associated with markers using forward and backward stepwise regression in mixed mode (Segura et al., 2012). All models incorporated marker-based kinship and population structure covariates as appropriate. Multiple testing correction was performed with a Bonferroni procedure (α = 0.05). Results were visualized with Manhattan plots (nominal visualization threshold set at −log10(p-value) = 5.5) and quantile–quantile (Q–Q) plots to assess test statistic inflation and model fit. LD was estimated by pairwise correlation coefficients (r²) to define candidate regions, and loci with −log10(p) > 3.6 (p < 2.51 × 10⁻⁴) were highlighted for further inspection. Multiple testing was addressed using the false discovery rate (FDR) method (Benjamini & Hochberg, 1995), and SNPs with q-values below 0.05 were considered significant.

### Genetic interactions analyses

To understand the genetic action of the candidate genes mapped in the GWAS we use a model based in Cockerham model (Kao & Zeng, 2002 & Cockerham, 1954) for detecting epistatic two-way bi-allelic interactions; denoting locus 1 with *A* and *a* alleles and locus 2 with *B* and *b* alleles. By doing this, we were able to test if the two markers interact each other. The statistical model was:

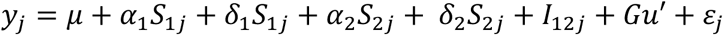

Where ***y***_***i***_ is the phenotypic value of clone *i*, ***μ*** is the overall mean, ***α***_**1**_ and ***α***_**2**_ are the additive effect of allele substitution at locus 1 and locus 2, respectively, ***δ***_**1**_ and ***δ***_**2**_ denote the corresponding dominance deviations, ***S***_**1*j***_ and ***S***_**2*j***_ represent the additive genotype coding for individual *j* at loci 1 and 2, respectively (coded as 0, 1, and 2 according to allele dosage). The terms ***αS***_**1*j***_ and ***αS***_**2*j***_ capture the linear effects of allele dosage at each locus, whereas ***δS***_**1*j***_ and ***δS***_**2*j***_ account for dominance deviations from the midpoint of the homozygotes., ***I***_**12*j***_ represents the epistatic interaction effects between loci 1 and loci 2 for individual *j* (additive x additive, additive x dominance, dominance x additive, dominance x dominance interactions), The kinship structure among individuals was modeled using the matrix ***Gu***′ and ***ε***_***i***_ is the residual error. A bonferroni threshold (∝ = 0.05) was used to correct the multiple comparisons. The proportion of phenotypic variance explained by the additive loci identified through GWAS, as well as by both additive and epistatic loci, was calculated as the r^2^ from a simple linear regression of these loci on mean trait performance.

### Candidate gene identification, functional annotation and haplotype block analysis

Candidate genes within a conservative window (±1 kb from each focal SNP based on LD results) were extracted and annotated using Gene Ontology (GO), InterPro and Pfam descriptors (El-Gebali et al., 2019). Homology and additional functional annotation were retrieved from National Centre for Biotechnology Information (NCBI) (http://www.ncbi.nlm.nih.gov/) and European Molecular Biology Laboratory European Bioinformatics Institute (EMBL-EBI) (https://www.ebi.ac.uk/) resources to support putative biological roles. Where appropriate, gene annotations were cross-referenced against published studies implicating pathways relevant to PPD. Candidate loci were prioritized based on statistical strength of association, biological plausibility and local LD patterns. Haplotype block analysis was conducted using *HaploView v.4.2* (Barrett et al., 2005), and gene structure visualizations were generated with *SnapGene v.7.0*. Haplotype blocks were defined based on linkage disequilibrium (LD) decay patterns inferred from the genome-wide association study (GWAS).

## Results

### Phenotypic variation and heritability of PPD

We evaluated phenotypic variation for PPD across multiple location–year combinations. Visual assessments of PPD ranged from 0.0 % to 94.73 % (mean = 31.78 %), while ImageJ-derived measurements ranged from 0.0 % to 99.92 % (mean = 30.62 %) with a coefficient of variation of 64.5 %). The Q–Q plots compare the observed residual distributions with the expected normal distributions for each trial or environment included in the PPD analyses. In most environments, the residuals are closely aligned with the reference line, suggesting that the residuals approximate a normal distribution and that the fitted models effectively capture the data’s variation (Supplementary Figure. S1). Broad-sense heritability estimated across environments varied between 0.22 and 0.58, with an overall mean of 0.48 (Supplementary Table S2), indicating a substantial genetic component to PPD expression alongside a large environmental contribution. Variance decomposition attributed 48 % of the phenotypic variance to genotypes × environments interaction (Vg×e) and 12 % to genotype main effects (VG), underscoring the importance of multi-environment evaluation in selection. The two scoring methods were highly concordant (correlation r = 0.97, Supplementary Figure. S2), supporting the adoption of the faster, visual assessments pipeline as a scalable alternative to ImageJ for routine breeding evaluations; until fully automated ImageJ workflows are implemented, visual scoring can continue as a practical interim solution.

### SNP distribution and linkage disequilibrium

After SNP calling and quality control, 27,047 high-quality biallelic SNPs were retained and mapped across the 18 cassava chromosomes (reported mapped span = 35 Mb). SNP marker distribution was heterogeneous across chromosomes: chromosome 1 carried the largest number of markers (n = 572) while chromosome 14 contained the fewest (n = 259) (Figure. 1a). Minor allele frequencies ranged from 0.05 to 0.50 with a mean MAF of 0.24 (Figure. 1b), and approximately 75 % of markers had allele frequencies exceeding 5 % in the evaluated panel, ensuring reasonable informativeness for association mapping.

**Figure 1.**
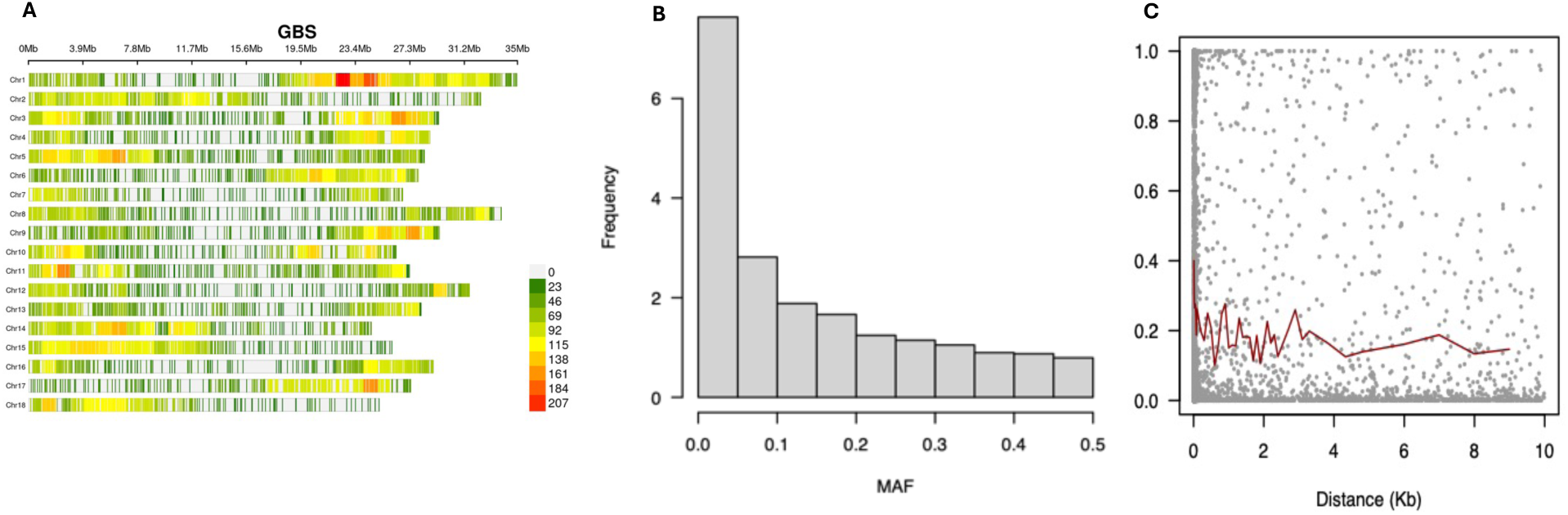
Overview of genotypic data: (**A)** SNP density distribution across the 18 chromosomes of cassava in the evaluated population**. (B)** Minor allele frequencies of the SNPs identified in the studied clones. (**C)** Linkage disequilibrium (LD) decay: This illustrates the squared correlation (r²) between markers and their physical distance in kilobases (kb).

We characterized linkage disequilibrium (LD) by computing pairwise r² across each chromosome to assess mapping resolution. Mean genome-wide r² was 0.19, with a peak LD near r² = 0.40 that decayed to r² ≈ 0.18 over short physical distances, on the order of ∼1 kb in the present panel (Figure. 1c). This rapid LD decay indicates fine local resolution for association mapping and guided the window sizes used to define candidate regions around significant markers.

### Population structure and relatedness

Population structure analyses were performed on the post-QC dataset used for structure inference (≈812 sample entries and ∼26k SNPs after the stricter QC used for these analyses). Principal component analysis (PCA) indicated modest stratification: the first two principal components (PC1 + PC2) together explained 14.23 % of the total genetic variance (Figure. 2a), and inclusion of PC3 increased explained variance to ∼17.69 % (Supplementary Figure. S3). Bayesian clustering and hierarchical analyses revealed two clear subgroups within the panel (Figure. 2b) that correspond broadly to historical germplasm and populations derived from recent genomic-selection cycles. Identity-by-state and kinship matrices showed generally low pairwise relatedness among most clones, consistent with a relatively diverse association panel and favorable conditions for detecting marker–trait associations with limited confounding by close kinship (Figure. 2b).

**Figure 2.**
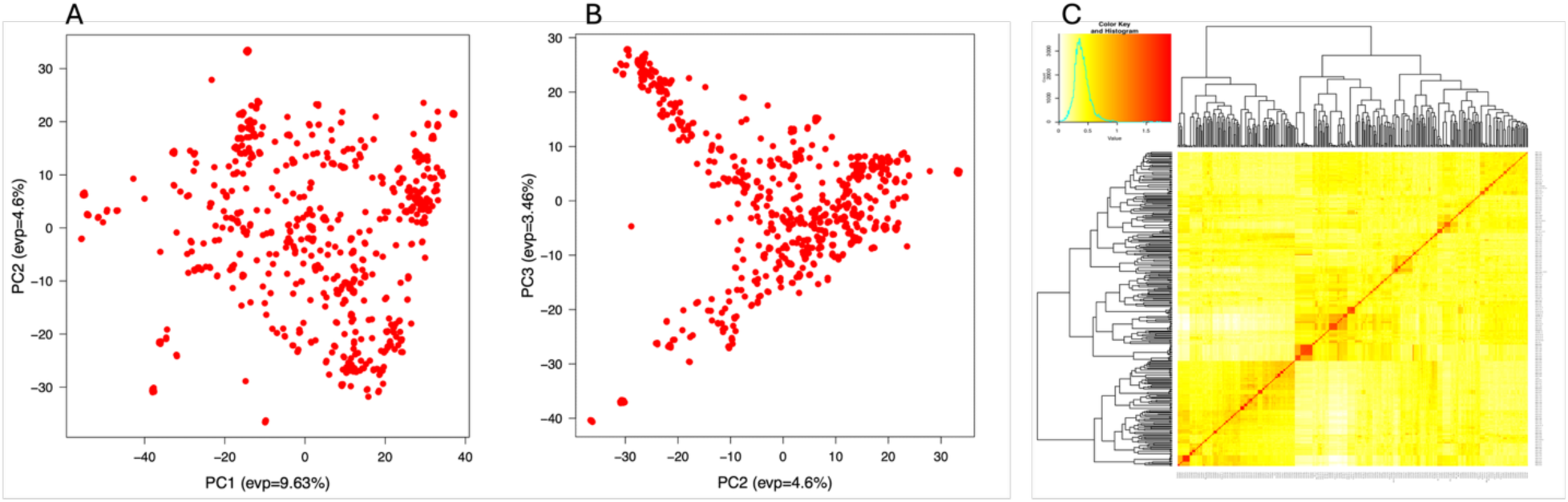
Population structure assessed by principal component analysis (PCA). **(A)** Scatter plot of the first and second principal components (PCs). **(B)** Scatter plot of the second and third principal components **(C)** Heatmap of the pairwise kinship matrix. Yellow and red represent weak and strong kinship relationships between genotypes, respectively. The clustering dendrogram is displayed outside the matrix.

### Genomic regions associated with post-harvest deterioration

We conducted GWAS using three complementary statistical models: MLMM, BLINK, and SUPER. The quantile–quantile (Q–Q) plots indicated minimal genomic inflation or deflation, with observed distributions closely matching expectations except for the significant associations that clearly deviated, confirming their statistical robustness (Figure. 3c, 3d). Across the three models, we identified 10 significant SNPs distributed on chromosomes 2, 5, and 13 (Supplementary Table S1; Figure. 3a, 3b). Notably, the markers located on chromosome 2 were consistently detected by all three models (MLMM, SUPER, BLINK) and were validated using both phenotyping approaches (visual scoring and ImageJ analysis). Two SNPs in this region, S2_11347402 (allele A/T) and S2_11347477 (allele T/G), together explained 8.34 % of the phenotypic variance (PVE) (Supplementary Table S1; Figure. 3a, 3b). This consistency across models and phenotyping methods highlights chromosome 2 as a key genomic region influencing PPD expression.

**Figure 3.**
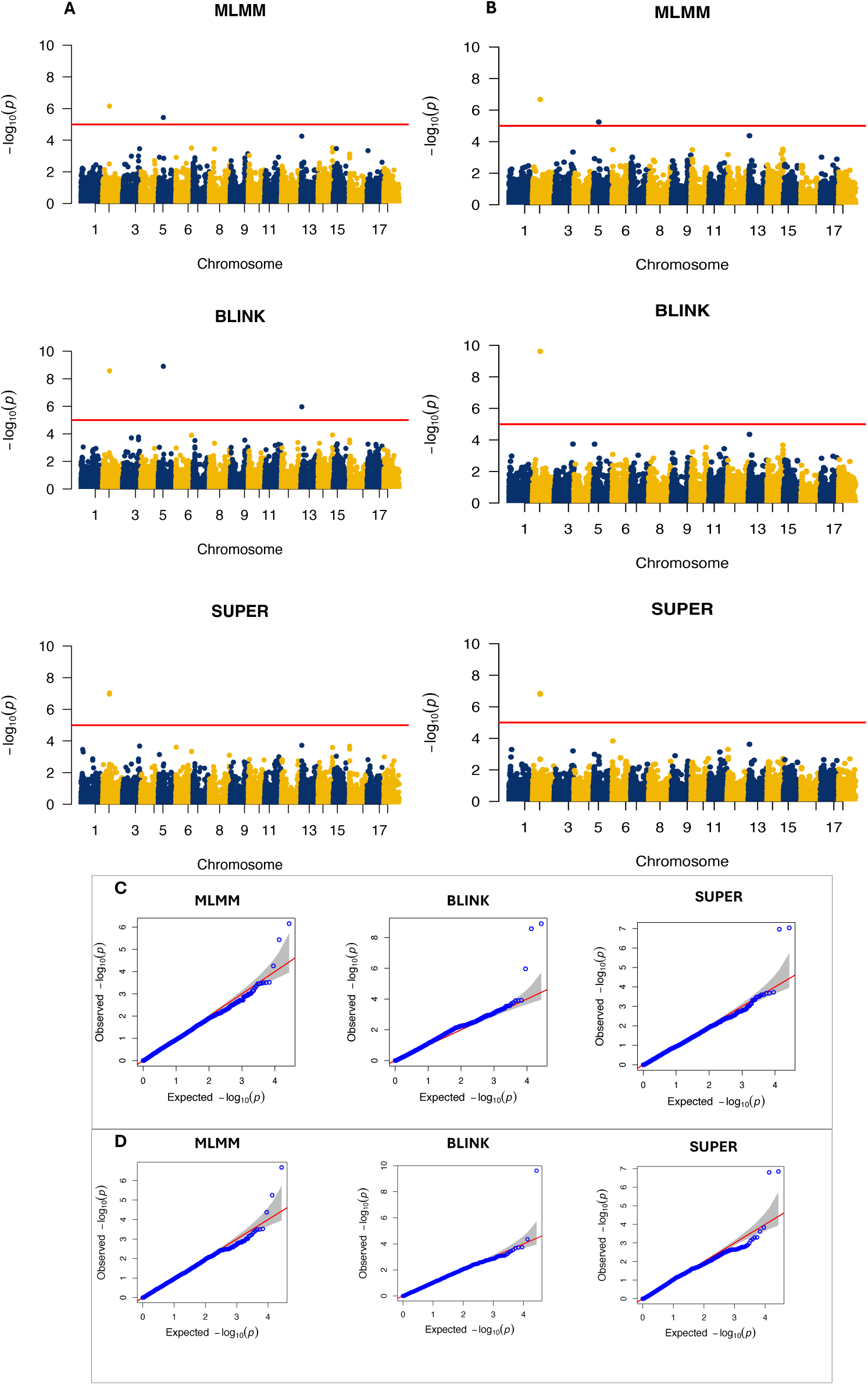
Results of genome-wide association studies (GWAS) using different statistical models. **(A)** Manhattan plots for PPD scored with ImageJ data employing MLMM, BLINK, and SUPER models. **(B)** Manhattan plots for PPD scored with visual data utilizing MLMM, BLINK, and SUPER models. **(C)** Quantile–quantile (Q–Q) plots for the models using ImageJ data. (D) Q–Q plots for the models using visual data.

The good agreement between the two phenotyping methods (visual and ImageJ) explains the consistency of the associations across analyses. On chromosome 5, a significant SNP (S5_ 8348709, allele A/G) was detected using the BLINK model with ImageJ data, explaining 23.87% of the phenotypic variance (PVE). On chromosome 13, the SNP S13_1760296 (alleles C/A) was identified by the BLINK model with ImageJ data, explaining 3.62 % of the PVE (Supplementary Table S1; Figure. 3a). In total the four significant SNPs detected on chromosomes 2, 5, and 13 accounted for 35.83 % of the total phenotypic variance in PPD observed across the population. Comprehensive details of the peak SNPs associated with all loci and their candidate genes are provided in the supplementary material (Supplementary Table S1).

### Prediction of candidate genes

Functional annotation of significant loci was performed using the cassava reference genome v6.1 available on the Phytozome platform (Goodstein et al., 2012). This analysis enabled the prediction of causal genes underlying loci significantly associated with PPD expression. In total, three candidate genes were identified across the significant SNPs detected by the different GWAS models. On chromosome 2, SNPs S2_11347402 and S2_11347477 were located within the coding region of the gene *Manes.02G152300*, which encodes an aquaporin transporter (PTHR19139//PTHR19139:SF145) (Figure. 4a). Aquaporins are transmembrane proteins that mediate the transport of water and small molecules, including glycerol, CO₂, hydrogen peroxide, and lactic acid, across biological membranes (Kapilan et al., 2018). Given their established roles in water and ROS transport, aquaporins are highly relevant to wound responses and early-stage PPD progression.

**Figure 4.**
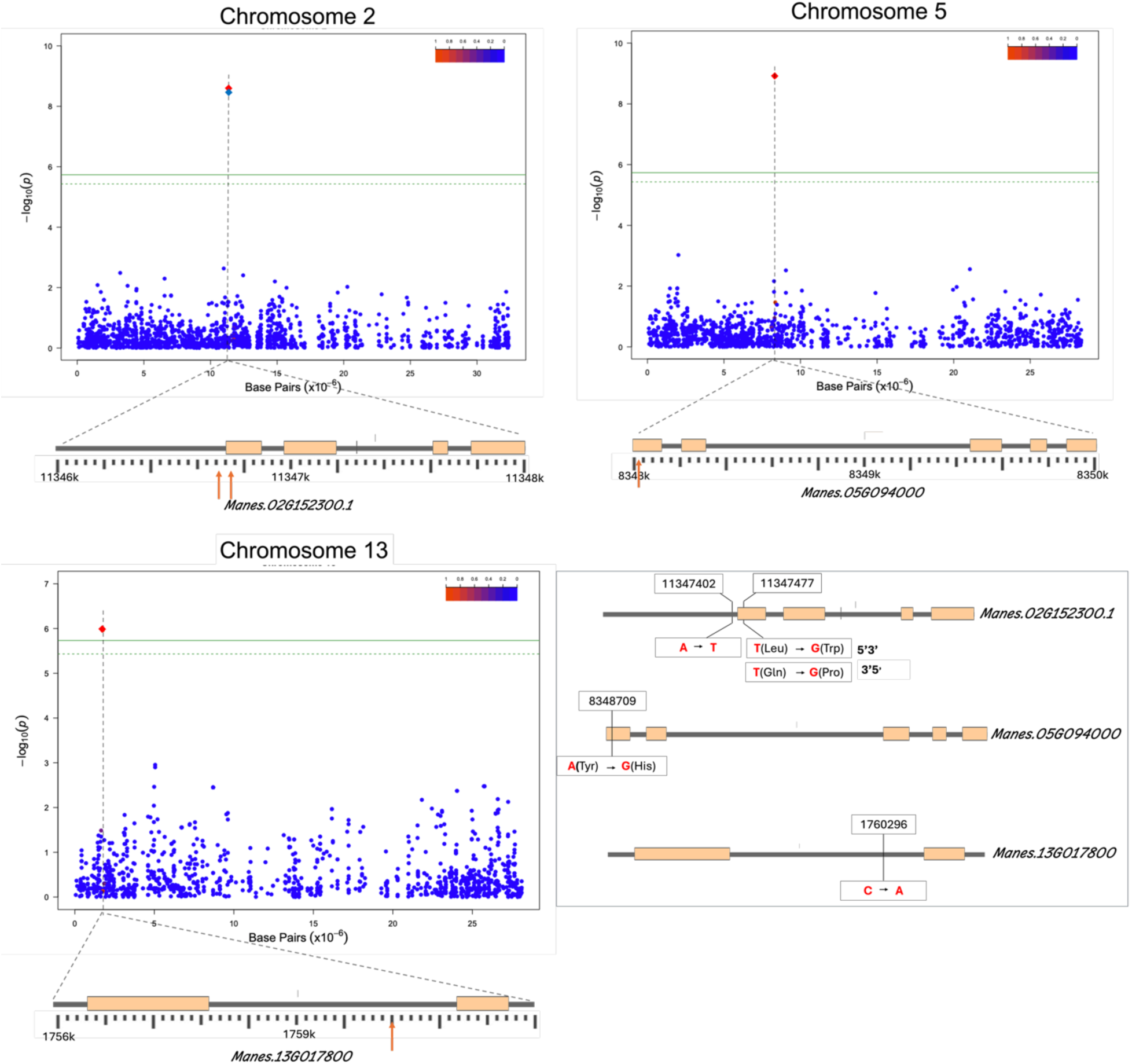
Candidate genes and local LD around peak SNPs. **(A–C)** Regional association plots for the three chromosomes showing –log10(p) of GWAS signals (top), pairwise LD (r²; color scale) and annotated genes in the interval (bottom). The lead SNP position is indicated by a dashed vertical line; the significance threshold is shown. Orange pointers mark SNP positions mapped to gene models. **(D)** Haplotype maps for candidate genes: coding regions (gray), coding sequences (bisque), mutated bases highlighted in red, and amino-acid changes (in parentheses) where applicable.

On chromosome 5, SNP S5_ 8348709 was located within the coding region of *Manes.05G094000*, which encodes a serine/threonine protein phosphatase (PTHR11668//PTHR1668:SF257) (Figure. 4b). These enzymes are central regulators of plant defensive responses to biotic and abiotic stress, often through the induction of oxidative stress and the generation of reactive oxygen species (ROS) that damage cellular structures. In chromosome 13, SNP S13_1760296 was positioned near *Manes.13G017800*, which encodes a leucine-rich repeat (LRR) protein kinase (PTHR27008//PTHR27008:SF4) (Figure. 4c). LRR kinases are essential in interpreting extracellular signals and initiating downstream transduction cascades.

Further inspection of allelic variants highlights the functional consequences of these SNPs. In *Manes.02G152300*, S2_11347477 results in a nonsynonymous substitution from leucine (Leu) to tryptophan (Trp), while S2_11347402 is located in the coding region. In Manes.05G094000, the SNP introduces an amino acid substitution from tyrosine (Tyr) to histidine (His), which could significantly affect protein folding and interactions, potentially explaining phenotypic differences between tolerant and susceptible clones. Finally, in *Manes.13G017800*, variation in the intergenic region likely influences transcriptional regulation of kinase activity, modulating the balance between ROS production and hypersensitive responses.

### Haplotype-based GWAS analysis

To characterize local haplotype structure around trait-associated loci, we examined four flanking SNPs upstream and downstream of each peak GWAS marker on chromosomes 2, 5 and 13, and defined haplotype blocks from local LD patterns. Mean haplotype-block sizes were ∼107 kb on chromosome 2, ∼47 kb on chromosome 5 and ∼202 kb on chromosome 13 (Figure. 5), reflecting chromosome-specific differences in local linkage disequilibrium and recombination. Block boundaries and local r² profiles informed the genomic intervals used for candidate-gene inspection. Haplotype composition was evaluated in a targeted subset of 19 clones selected to span the phenotypic range (low, medium and high PPD). Among the low-PPD group, 9 accessions carried one or two copies of the chromosome-2 alleles (S2_11347402 / S2_11347477) but carried zero copies of the variant alleles at the chromosome-5 and chromosome-13 loci. In contrast, clones with medium and high PPD typically carried fewer or no copies of the protective chromosome-2 alleles while frequently carrying two copies of the variant alleles on chromosomes 5 and 13. Boxplot summaries (Figure. 5) show a clear stepwise increase in PPD severity associated with presence of the derived alleles on chromosomes 5 and 13, supporting a contributory role of these loci to symptom severity. Collectively, these haplotype patterns are consistent with a multi-locus genetic architecture in which alleles on chromosomes 5 and 13 increase susceptibility, whereas alleles on chromosome 2 are associated with reduced PPD. These results complement the single-marker GWAS by (i) refining the physical intervals that harbor candidate genes, (ii) revealing haplotypic backgrounds that differentiate tolerant and susceptible clones, and (iii) suggesting that favorable allele combinations at chromosome-2 loci may partially mitigate the adverse effects of susceptibility alleles on chromosomes 5 and 13.

**Figure 5.**
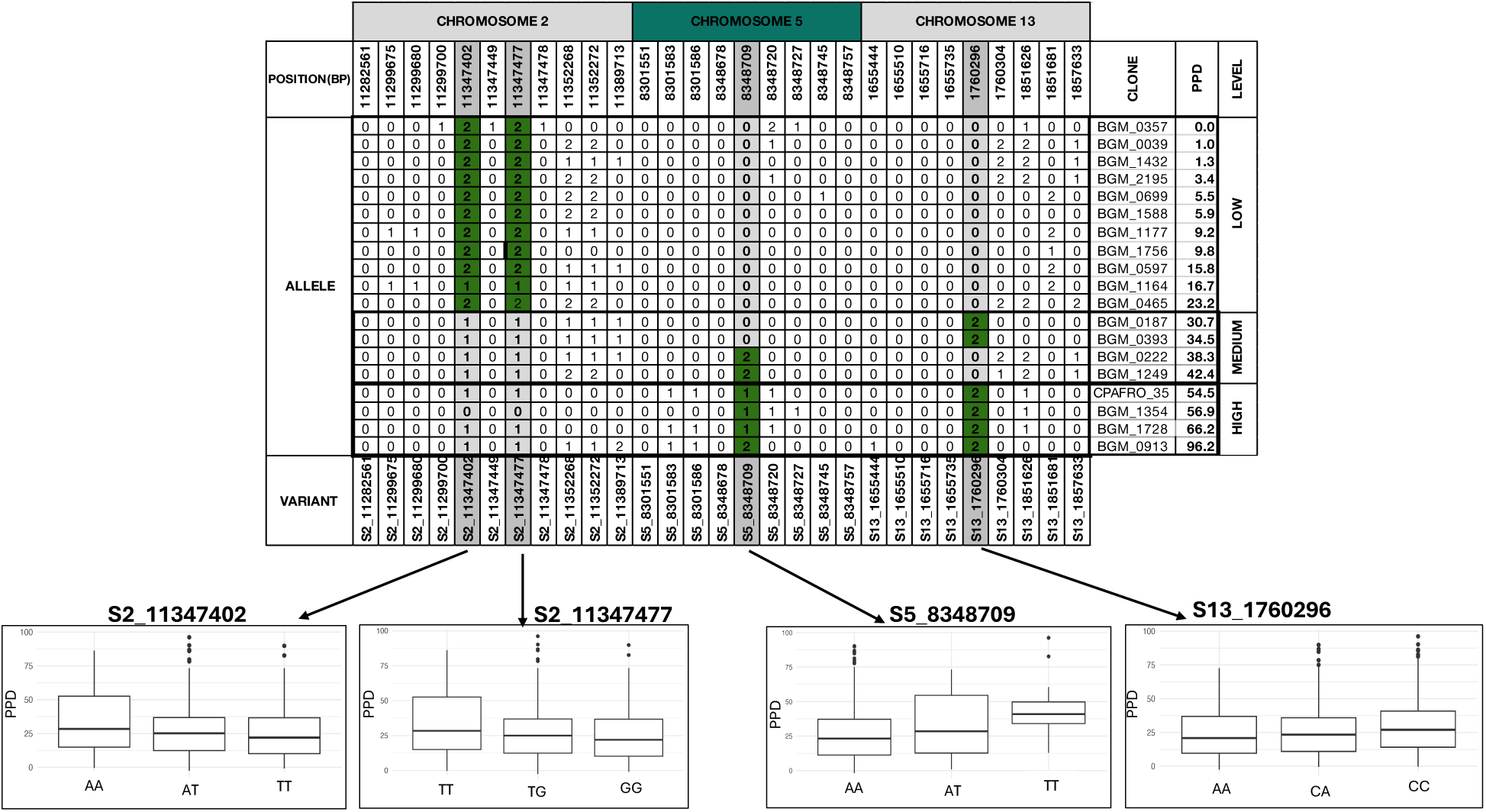
Haplotype analysis of candidate genes involved in PPD expression across the cassava genome. At the top, the SNP analysis is presented for chromosomes two, five, and thirteen, with the corresponding positions of the variants indicated in gray. On the right, the clones are categorized into three groups based on the severity of PPD symptoms: low, medium, and high. The numbers zero, one, and two represent the number of SNP copies present in each genotype. In the bottom section, a boxplot illustrates the levels of PPD symptoms associated with each genotypic allele combination, showing data for those with zero, one or two copies of the reference SNPs.

### Statistical inference of epistatic interactions among candidate genes

The analysis of genetic interactions among candidate genes provides deeper insights into the genetic basis of complex traits such as PPD and facilitates the identification of favorable allelic combinations for pyramiding in breeding strategies. In this study, we focused on alleles located on chromosomes 2, 5, and 13, as revealed by GWAS, and evaluated their potential interactions through both haplotype and epistasis analyses. Haplotype analysis suggested that specific allele combinations can attenuate the impact of metabolic pathways underlying PPD expression in cassava roots. To quantify these effects, we performed an epistasis analysis using four SNPs, arranged into pairs. The results revealed clear interaction patterns: 1) The allele combination S2_11347477_GG × S5_8348709_AA was associated with significantly reduced PPD expression, consistent with an additive × additive interaction, with a estimated value of –43.25; 2) In contrast, the combination S5_ 8348709_AT × S13_1760296_CC was linked to markedly higher PPD levels, showing a dominance × additive effect, with value of 52.25 increasing of PPD expression in the genotypes with this allele combination related with the population mean; 3) Additionally, a significant additive × dominance interaction was detected between S2_11347477_GG × S13_1760296_CA, which reduced PPD expression by –26.25 compared with the population mean. These results (Supplementary Table S3; Figure. 6) highlight the importance of epistatic interactions in shaping PPD expression, emphasizing that tolerance is not solely determined by individual loci but also by the interaction among alleles across different genomic regions.

**Figure 6.**
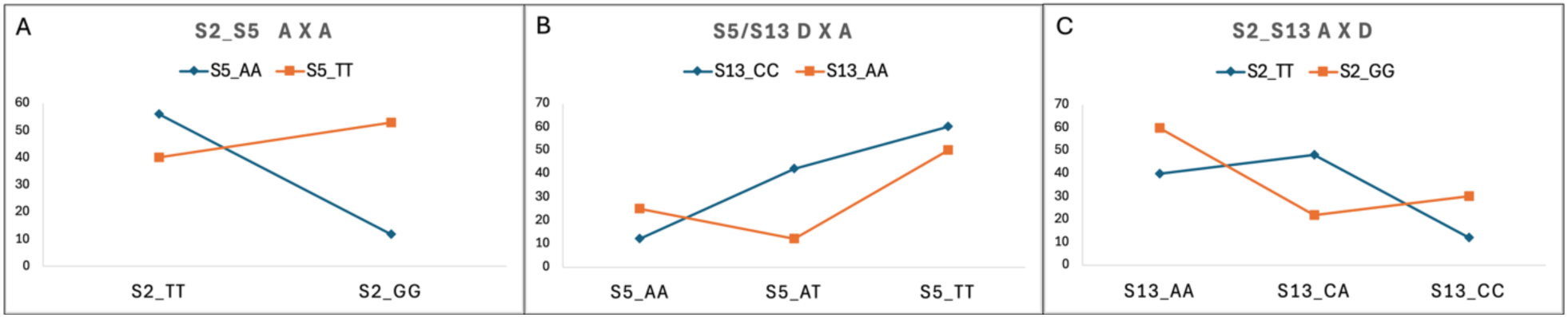
Epistasis plots for SNP pairs. **(A)** Additive × additive interaction between S2_11347477 and S5_ 8348709; the x-axis represent the genotypes at SNP S2_11347477 (TT/GG), the colored lines correspond to genotypes classes at SNP S5_ 8348709 (AA blue and TT orange) and the y-axis represents the phenotypic value of the PPD; **(B)** dominance × additive interaction between S5_ 8348709 and S13_1760296; the x-axis represent the genotypes at SNP S5_ 8348709 (AA/AT/TT), the colored lines correspond to genotypes classes at SNP S13_1760296 (CC blue and AA orange) and the y-axis represents the phenotypic value of the PPD **(C)** additive × dominance interaction between S2_11347477 and S13_1760296. the x-axis represents the genotypes at SNP S13_1760296 (AA/CA/CC), the colored lines correspond to genotypes classes at SNP S2_11347477 (TT Blue and GG orange) and the y-axis represents the phenotypic value of the PPD.

## Discussion

### Genomic approaches and the challenge of studying PPD

Genomics-driven approaches have become indispensable to modern crop improvement because they accelerate the development of varieties that combine favorable traits with stability across environments. For complex physiological traits such as PPD in cassava, understanding the underlying genetic architecture is a prerequisite for efficient selection. PPD is an intrinsically complex phenotype: it is triggered by harvest-induced wounding and involves coordinated responses across stress signaling, ROS dynamics, water and solute flux, and multiple metabolic pathways that together determine whether a genotype will be highly susceptible or comparatively tolerant.

Our findings sit within a growing literature that has characterized biochemical, transcriptomic and genetic contributors to PPD, including metabolomic studies (Lebot et al., 2023), transgenic experiments (Ma, et al., 2022), surveys of Brazilian germplasm (Venturini et al., 2016), and transcriptome profiling (Reilly et al., 2007). Integrating these layers (genome, transcriptome, and metabolome) strengthens candidate gene prioritization because it links statistical association to plausible biological mechanisms. A major strength of this study is the use of multi-environment, multi-year phenotyping, which increases power to detect reproducible associations and helps to partition genetic signal from environmentally driven variation.

### Panel diversity, structure, and heritability

The evaluation panel showed wide phenotypic variation for PPD (0–94 %), indicating segregation of alleles with contrasting effects. The panel’s average MAF = 24 % provides reasonable power for association detection while highlighting that some alleles may be moderately rare and therefore could have limited transferability across germplasm sets. Principal component analysis separated samples into two clusters but did not indicate strong stratification that would confound GWAS. Kinship estimates were low, consistent with considerable diversity in the panel, a favorable scenario for association mapping. Heritability estimates varied among environments with a mean near 48 %, which aligns well with Venturini et al. (2016), who reported ∼52 %. An intermediate heritability implies that genetic improvement is attainable but will be moderated by environmental influence and G×E; thus, breeding strategies must account for both genetic effects and environmental robustness.

### GWAS strategy and functional annotation

To maximize detection and to reduce false positives, we applied three complementary models: MLMM, BLINK and SUPER. Each model balances control of structure/kinship and computational efficiency differently; convergence of signals across models increases confidence in putative loci. MLMM was particularly sensitive to signals on chromosome 2, BLINK captured associations across chromosomes 2, 5 and 13, and SUPER both confirmed chromosome 2 signals and revealed additional local variants. All three models consistently identified S2_11347402 as a high-confidence SNP associated with PPD.

This study identifies three key biological and biochemical processes that occur after the harvest process and are associated with the candidate genes proposed for the early stages of PPD expression. The first process involves aquaporins, wich facilitate transmembrane water and small-molecule movement and are known to respond rapidly to wounding and osmotic changes. Reilly et al. (2007) reported early upregulation of upregulation of Plasma membrane Intrinsic Protein PIP1 and PIP2 family members during PPD; Zainuddin et al. (2018) also highlight the importance of water/ion transport in the early wound response. We provided two SNPs on chromosome 2 (S2_11347402 and S2_11347477) jointly explain ∼8.9 % of phenotypic variance. They map to a region that includes *Manes.02G152300*, annotated as an aquaporin (PIP-type) gene. We detected a coding-frame mutation co-located with S2_11347402 that results in a notable amino acid change between leucine (Leu) and tryptophan (Trp) (Figure 4d). The physicochemical change (hydrophobic aliphatic residue to bulkier aromatic residue) could alter channel conformation, gating dynamics, or interactions with regulatory partners, consistent with the observation that clones homozygous for the alternative allele show improved tolerance (Figure 5). These lines of evidence support a functional role for this aquaporin in modulating early PPD dynamics.

In following biochemical process related to PPD, Serine/threonine phosphatases are central regulators of signal transduction, cellular metabolism, and both biotic and abiotic stress responses (Máthé et al., 2019) the context of PPD, these enzymes influence oxidative stress pathways and the generation of reactive oxygen species (ROS), which underlie the characteristic darkened tissue observed in advanced PPD (García et al., 2013).

In biochemical processes associated with PPD, serine/threonine phosphatases act as central regulators of signal transduction, cellular metabolism, and responses to both biotic and abiotic stresses (Máthé et al., 2019). In this context, these enzymes are thought to modulate oxidative stress pathways and the production of reactive oxygen species (ROS), which contribute to the characteristic tissue darkening observed during advanced stages of PPD (García et al., 2013).They also participate in signaling cascades involving phytohormones such as jasmonic acid. Pharmacological studies using phosphatase inhibitors (e.g., okadaic acid) further underscore the role of phosphatase activity in modulating stress responses. (Choudhury et al., 2017) our findings reveal in chromosome 5, the SNP S5_8348709 explains approximately 23.87 % of the phenotypic variation in our panel. This variant maps to *Manes.05G094000*, which encodes a serine/threonine phosphatase. In our dataset, Figure 4d shows a non-synonymous substitution at this locus of tyrosine (Tyr) to histidine (His), a change in residue chemistry that may affect local folding, charge distribution, or protein–protein interactions. Figure 5 demonstrates that clones carrying the alternative allele display a marked increase in PPD. Together, these lines of evidence support a plausible functional role for this phosphatase in modulating PPD progression.

One of the late symptoms observed in the expression of PPD is the appearance of blue-dark marks in the tissues, which result from hypersensitivity and cell death. In this process of activation leucine-rich repeat receptor-like kinase LRR-RLKs play a crucial role in the membrane-localized receptors that mediate extracellular signal recognition and downstream signal transduction, and they play central roles in responses to both biotic and abiotic stress. of these receptors are frequently associated with triggered immunity (TI), a response that includes rapid ROS accumulation and a cascade of signaling events that may culminate in hypersensitive responses and programmed cell death (Ma, Hu, et al., 2022). Such immune-related responses are consistent with the late stages of PPD, when tissue integrity is severely compromised and necrotic regions become prevalent (Djabou et al., 2017). In our findings we report on chromosome 13 explains approximately 3.62 % of the phenotypic variation, the SNP S13_1760296. This variant lies in *Manes.13G017800*, which encodes a (LRR-RLK). This association is reflected in Figure 5, where clones carrying the alternative allele show higher PPD scores. Thus, one plausible hypothesis is that the variant at S13_1760296 predisposes roots to an exaggerated ROS-mediated hypersensitivity response after harvest, accelerating tissue collapse.

Understanding epistatic interactions is essential for analyzing the interactions between candidate genes and optimizing the pyramiding of genes in future genomic selection processes. Our epistasis tests revealed significant interactions between loci on chromosomes 2 and 5; when favorable alleles are combined at these loci, the average PPD score is reduced by 43.25 units (on the phenotyping scale used). In this study, we provide evidence of a significant interaction between the transcript products of genes *Manes.02G152300* and *Manes.05G094000*. The first gene encodes for aquaporins, which have been identified as essential regulators in molecular transport and play a crucial role in the plant’s stress response. In the context of PPD, this means that if a plant possesses a favorable allele combination in its genotype, it will exhibit improved mechanisms to control the PPD reaction during the initial stages or hours. This finding is linked to the second gene, responsible for transcribing antioxidant molecules. When a plant possesses a favorable combination of alleles for certain genes, it can produce a regulated amount of molecules involved in the oxidative process. With effective transport, this enables the creation of a complex metabolic pathway, allowing tissue cells to mount a slower but more effective stress response after damage occurs during harvesting or transportation. Importantly, our statistical analysis provides strong evidence of this epistatic interaction, which can be demonstrated in the phenotypic expression of genotypes that harbor the genetic combinations for both genes. As a result, these plants display long-shelf life and low symptoms of PPD. This information provides an excellent opportunity to utilize the most favorable allele combinations across different loci, aiming for optimal effects in the pathways associated with PPD’s early and late symptoms. This approach will be valuable for developing the new genomic selection population in the coming years.

Together, the statistical and functional evidence point to four SNPs distributed across three chromosomes that jointly explain 35.83 % of the phenotypic variance. Each SNP maps within or near genes with clear roles in the physiological processes triggered by harvest wounding and mechanical injury. From these data we can reasonably hypothesize that there is a genetic interaction among the various regions where the SNPs are located, as well as interactions between the resulting products of the metabolic pathways during both the early and late phases of post-harvest physiological deterioration.

### Perspectives for cassava breeding

The present study provides a comprehensive catalog of loci that significantly affect PPD expression. By utilizing historical data from multi-environment evaluations and employing two assessment methods, visual analysis and image, along with high-density genome-wide SNP data from GBS, we achieved a higher mapping resolution. As a result, we can identify candidate genes closely related to the PPD biological processes that occur in the roots after harvesting. We also identified SNP variants that are advantageous for extending cassava’s shelf life. Furthermore, this study provides specific allele combinations in the haplotype. If this information could be converted into allele-specific high-throughput SNP markers, it would enable fast and efficient screening and identification of individuals in the segregating population that carry the favorable alleles during the genomic selection process. Additionally, our study offers substantial insights into the epistatic interactions between genes on chromosomes 2 and 5. This information enhances our understanding of the strategies to be used in breeding programs. By recognizing the strong relationship between the transcripts of *Manes.02G152300* and *Manes.05G094000*, we can utilize this information to select favorable alleles in the segregating population, facilitating the pyramiding of these genes in future generations.

Nevertheless, this information is derived from a rigorous statistical analysis and data sourced from a GenBank on genes and protein information. Our findings have the potential to advance future functional studies that enhance our understanding of the interactions among candidate genes. Furthermore, to explore how these genes’ transcriptional products and their corresponding proteins interact to regulate the processes affecting root tissue during the hours following harvest and under transportation stress. Additionally, it would be interesting to develop CRISPR-Cas9 methodologies to silence gene expression. This could be validated with biological evidence demonstrating that these genes play a crucial role in the expression of PPD. The studies mentioned, along with other related research, would provide valuable insights into the pathways involved in PPD expression. This information would be essential for the cassava community, aiding in the management of breeding processes through molecular techniques, biochemical approaches, genomic selection, and technology transfer to producers.

### Concluding remarks

In this study, we demonstrate the value of integrating a historical, multi-year phenotypic dataset with high-density SNP genotyping and complementary GWAS models to identify genomic regions associated with delayed PPD in cassava. This combined methodological framework increases power to detect reproducible associations and accelerates the discovery of candidate variants that can be translated into breeding tools for complex, environmentally sensitive traits. We identified four SNPs across three chromosomes that together explain 35.83 % of the phenotypic variance for PPD in the Brazilian germplasm panel. These loci are associated with three biologically plausible candidate genes: an aquaporin (*Manes.02G152300*), a serine/threonine phosphatase (*Manes.05G094000*), and an LRR-RLK (*Manes.13G017800*). Each gene is linked to processes relevant to early (water and solute transport, membrane integrity) and late (ROS-mediated signaling, hypersensitivity) stages of PPD. Significant epistatic interactions were also detected, particularly between loci on chromosomes 2 and 5.

While these results provide substantial insights into the genetic architecture of PPD, it is important to recognize additional factors that may influence trait expression. Heritability and environmental effects remain critical considerations, as they directly affect the stability of PPD-related traits across diverse production environments. Likewise, the role of heterozygosity within the population warrants attention, since it may shape both allele effects and the efficiency of selection strategies aimed at reducing PPD. Even with these complexities, the loci and candidate genes identified here hold strong potential for integration into MAS and GS pipelines, offering practical tools for breeding programs. We anticipate that these findings will provide a valuable resource for the cassava research community and, in the long term, contribute to enhancing the economic viability and sustainability of cassava production for farmers, processors, and retailers alike.

