## Supplementary figures and images for "Unraveling candidate genomic regions responsible for delayed post-harvest deterioration in Cassava (*Manihot esculenta* Crantz)"

### Supplementary_Figure_S1

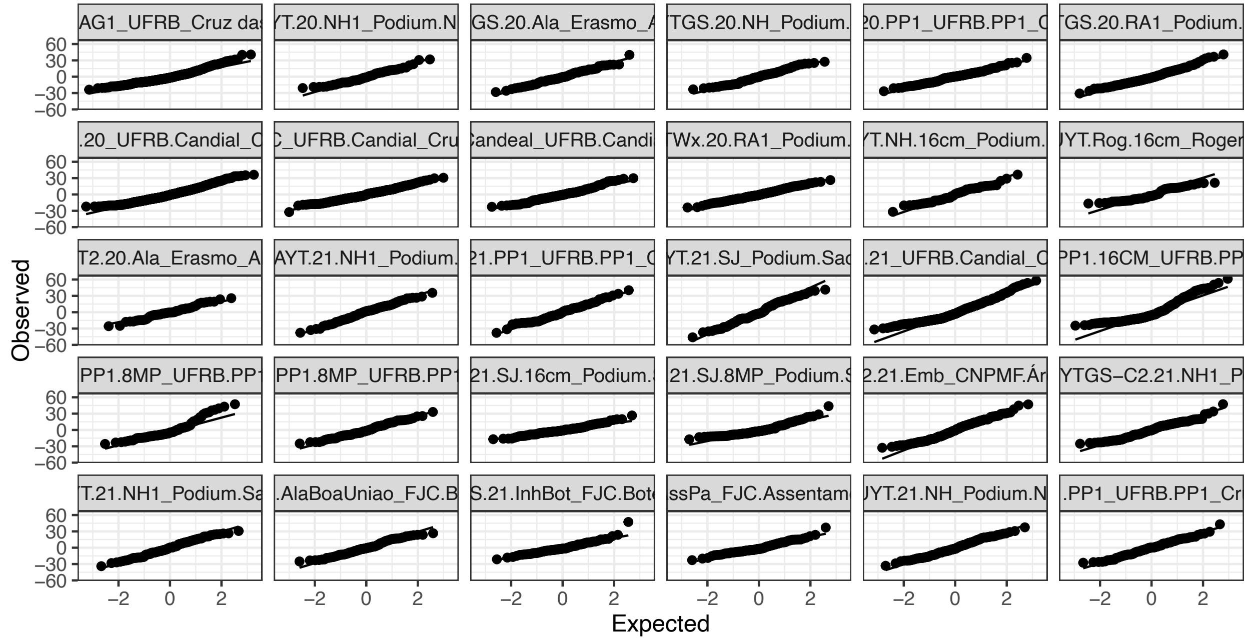

### Supplementary_Figure_S2

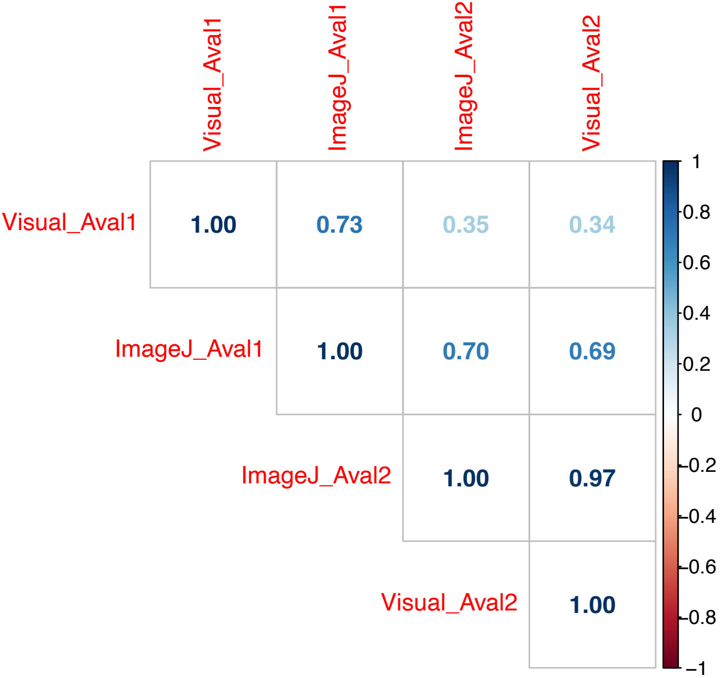

### Supplementary_Figure_S3

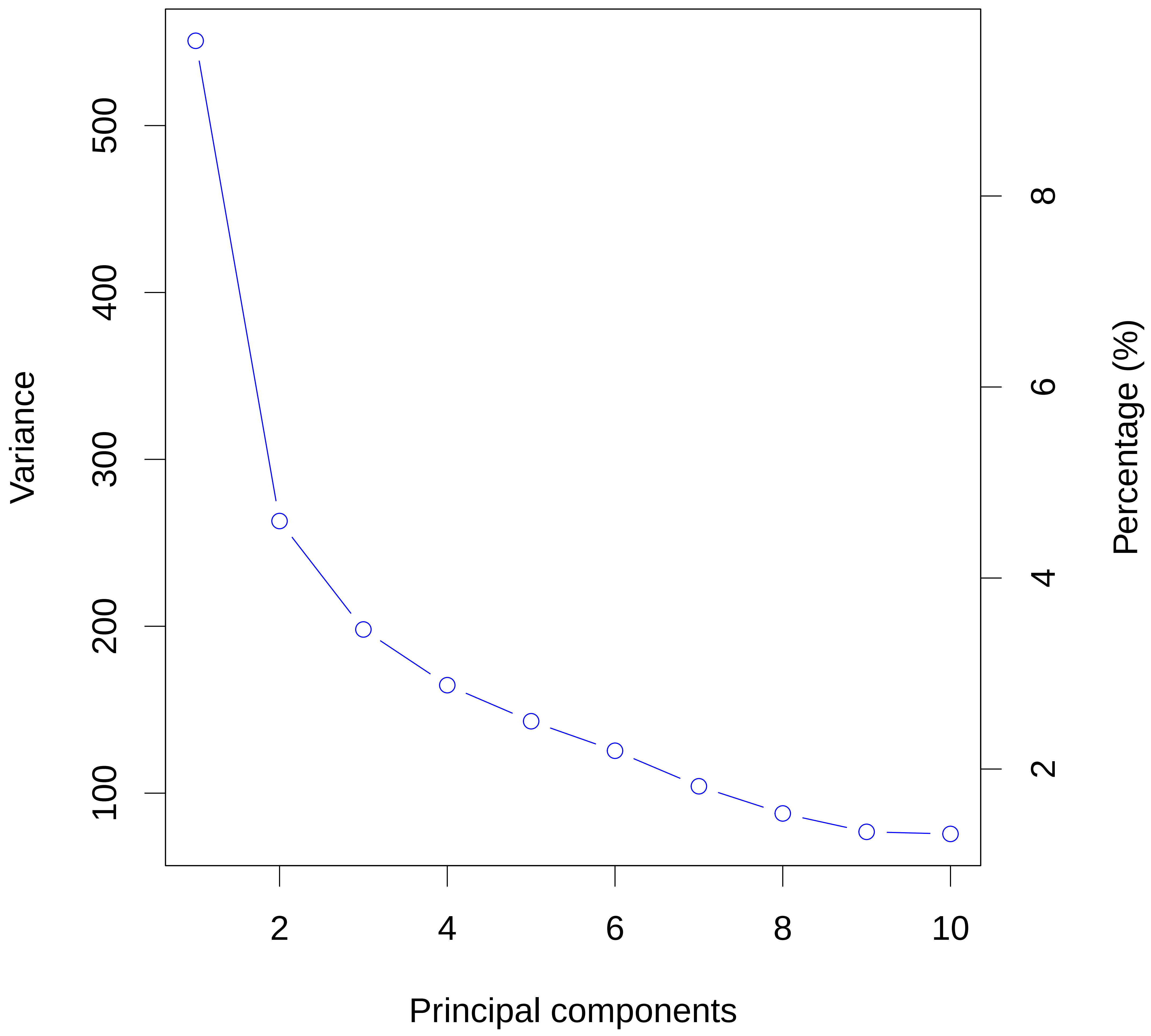
